# Zfp750 prevents oral adhesions and promotes temporary epithelial fusions

**DOI:** 10.64898/2026.02.12.705205

**Authors:** S Singh, E Adelizzi, C Heffner, S Curtis, K Duncan, W Awotoye, J Olotu, T Busch, WL Adeyemo, L JJ Gowans, T Naicker, SA Murray, A Butali, EJ Leslie-Clarkson, M Dunnwald, RA Cornell

**Affiliations:** Department of Oral Health Sciences, School of Dentistry, University of Washington, Seattle, WA; Department of Anatomy and Cell Biology, University of Iowa, Iowa City, IA; The Jackson Laboratory, Bar Harbor, ME; Department of Human Genetics, Emory University School of Medicine, Atlanta, GA; Department of Oral Pathology, Radiology and Medicine, College of Dentistry, University of Iowa, Iowa City, IA; University of Lagos, Lagos, Nigeria; Kwame Nkurumah University of Science and Technology, Kumasi, Ghana; Kwazulu Natal University, Durban, South Africa; Interdisciplinary Graduate Program in Genetics, University of Iowa, Iowa City, IA

**Keywords:** periderm, *ZNF750*, cleft palate, development

## Abstract

The differentiation cascade that converts basal keratinocytes into suprabasal layers, including periderm, depends on the activity of transcription factors. Mutations in the genes encoding many of these transcription factors, including *TP63, IRF6* and *GRHL3*, disrupt periderm development. Such mutations can also interfere with embryonic fusion and septation events that depend on periderm development, including palatogenesis, digit separation and the formation of temporary epithelial fusions between digits, between eyelids, and between pinnae and the scalp. ZNF750 (Zfp750 in the mouse) is a transcription factor required for keratinocyte differentiation, but whether mutations in *ZNF750* contribute risk for orofacial cleft, and the role of Zfp750 in periderm development, are unknown. To address these questions we sequenced *ZNF750* in 5,659 individuals including 2,125 with nonsyndromic OFC. We identify 33 rare missense variants with frequencies less than 0.1% in gnomAD. Of these, about half are predicted to be damaging with *in silico* tools. Collectively, these missense variants are not overtransmitted from parents to children with OFCs. Two of the variants have lower activity than the reference variant in a zebrafish embryo-based assay but no phenotype in the corresponding murine model. However, in murine embryos homozygous for a frame-shift mutation in *Zfp750* (*Zfp750*^*fs*^) that we generated, palatal shelves are fused but intra-oral adhesions are present, a phenotype seen in murine mutants of several *bonafide* OFC genes. In addition, temporary epithelial fusions are absent in *Zfp750*^*fs*^ neonates. RNA sequencing of forelimbs from *Zfp750*^*fs*^ embryos reveals decreased expression of epidermal terminal differentiation genes, and both increased and decreased expression of distinct periderm genes. Immunofluorescence shows the consistent presence of periderm proteins within the oral adhesions in *Zfp750*^*fs/fs*^ embryos. Together these studies suggest that while mutations in *ZNF750* are not a major contributor to OFC risk, Zfp750 does contribute to periderm-dependent morphogenic events.

## Introduction

The initial stratification of ectoderm-derived epithelial cells yields the periderm, a simple squamous epithelium covering the developing epidermis and oral epithelium until underlying epithelial cells are fully functional (Hammond et al. 2019; Holbrook and Odland 1975). In the oral cavity, periderm prevents inappropriate fusion between two adjacent epithelia during embryogenesis (Richardson et al. 2014). Consequently, the dissolution of the periderm at the medial edge is necessary for the two palatal shelves to fuse (Bush and Jiang 2012). During this process, periderm cells partially dedifferentiate and migrate toward the oral and nasal sides of the palatal shelf (Saroya et al. 2023; Teng et al. 2022).

Variants in genes encoding members of the transcriptional regulatory network governing the fate and differentiation of ectoderm-derived cells cause a spectrum of syndromes, known as peridermopathies, characterized by features such as orofacial cleft (OFC), digit anomalies, and epithelial webbing. Notable among these syndromes is Van der Woude syndrome (Kondo et al. 2002; Peyrard-Janvid et al. 2014; Robinson et al. 2025), the most common syndrome that includes OFC. Heterozygous variants in three genes (*IRF6, GRHL3, PRKCI*), all members of a regulatory network governing periderm development, cause this syndrome and common variants near *IRF6* and *GRHL3* are also associated with nonsyndromic OFC (nsOFC) (Beaty et al. 2010; Leslie et al. 2017; Leslie et al. 2016; Zucchero et al. 2004). Variants in *TP63* can cause several ectodermal dysplasia syndromes than can include OFC (OMIM:603273) and TP63 and IRF6 activate the expression of *IRF6* and *GRHL3*, respectively (de la Garza et al.2013; Moretti et al. 2010). Therefore, because mutations in genes encoding members of a regulatory network governing epithelial differentiation can cause OFC, other genes in this network are candidates to harbor mutations with this effect.

*ZNF750* encodes a transcription factor that regulates epidermal differentiation (Cohen et al. 2012; Sen et al. 2012) and is highly expressed in epithelial cells, including in the periderm (Li et al. 2019). A heterozygous frameshift mutation in *ZNF750* was found to perfectly segregate with affected individuals in a large pedigree with seborrhea-like dermatitis with psoriasiform elements (Birnbaum et al. 2006). Rare promoter variants of *ZNF750* are also associated with psoriasis phenotypes (Birnbaum et al. 2011). In mice, constitutive and epidermal-specific deletion of *Zfp750* leads to epidermal differentiation defects (Butera et al. 2023; Schwartz et al. 2024). Although the periderm was not a focus of either of the aforementioned studies, in both types of knockout embryos eyes were open at E16.5. Eyelids are typically fused at birth in wildtypes; transient epithelial fusions including between eyelids are accompanied by an accumulation of periderm cells (Maconnachie 1979). In the epidermal-specific *Zfp750* knockout embryos, palatal shelves were fused, suggesting Zfp750 is dispensable for palatogenesis. However, this issue is unresolved because the *Krt14* promoter driving Cre-recombinase in this model is active after the periderm has become a distinct layer (Rhea et al. 2025).

Here we sequence *ZNF750* exons in patients with nsOFC and assess whether coding variants detected in them retain function in zebrafish and mouse embryo-based assays. We also generate embryos homozygous for a frameshift mutation in *Zfp750* and assess them for oral adhesions and the absence of transient epithelial fusions, both attributed to abnormal periderm.

## Results

### ZNF750 variants predicted to be damaging are detected in individuals with nsOFC

To address whether variants in *ZNF750* could increase risk for OFCs we sequenced *ZNF750* exons in 2,125 individuals with nsOFC and their parents (when available), totalling 5,659 individuals (Fig. 1A). We identified 33 missense variants that were either absent from the Genome Aggregation Database or present at frequencies less than 0.1% (Appendix Table 1). Of the 33 missense variants, 18 were predicted to be damaging using *in silico* prediction tools. Notably, we did not identify any variants predicted to be loss of function (i.e., frameshift, nonsense, or splice variants) nor did we identify any variants that occurred *de novo*. In addition, of the variants meeting our criteria for allele frequency, impact, and *in silico* predictions, half were found only in unaffected parents and were not transmitted. Variants were distributed throughout the protein sequence but were not found within the zinc-finger domain (Fig. 1A). We statistically tested transmission in 759 complete trios (child with OFC and both parents) using a transmission disequilibrium test but found no evidence of over-transmission (*P* = 0.67).

**Figure 1:**
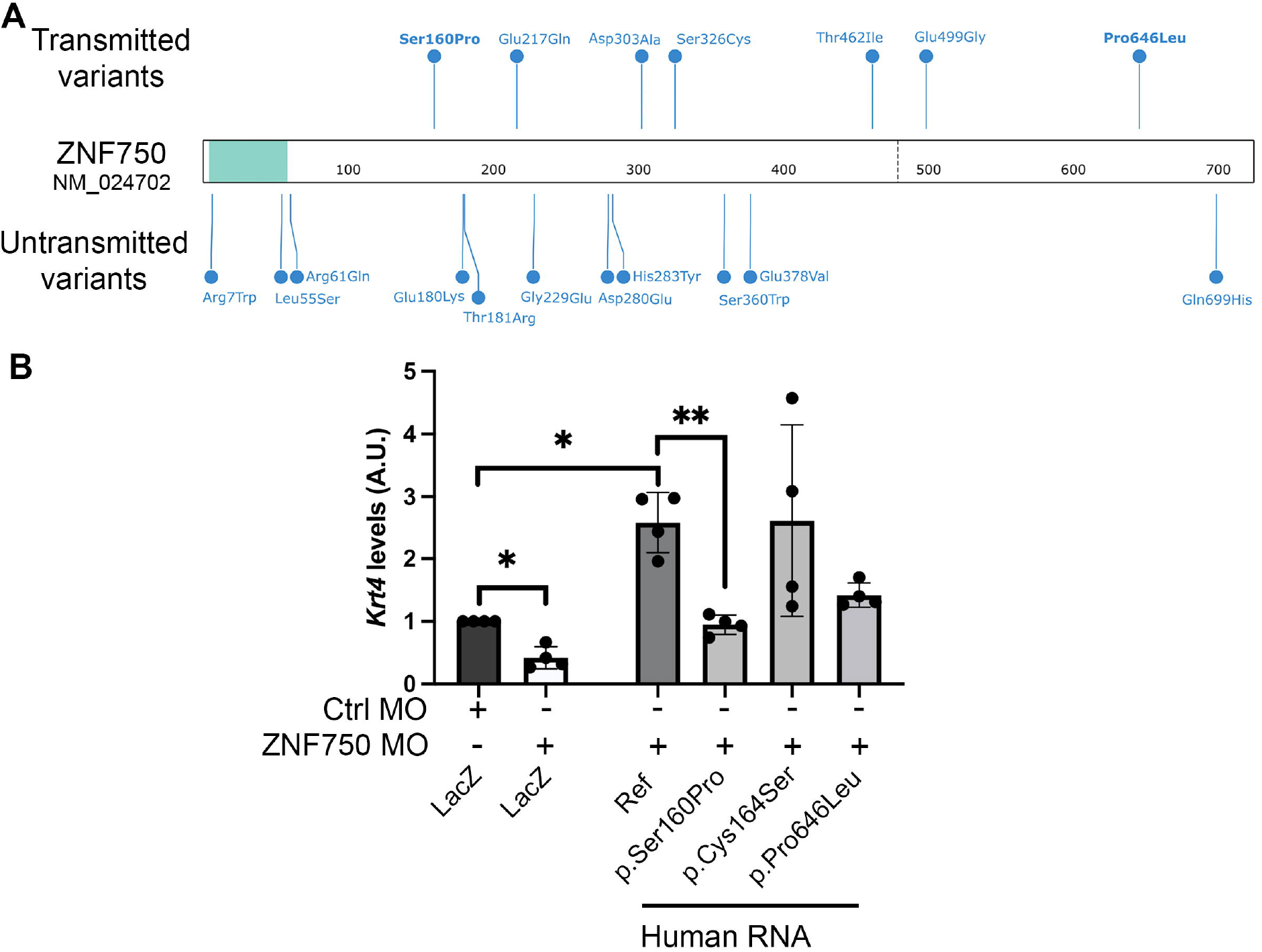
Candidate *ZNF750* variants do not affect periderm differentiation in zebrafish functional assays. (**A**) Rare *ZNF50* variants predicted to be damaging with *in silico* prediction tools. Variants transmitted to child with OFC are on top. Variants found in unaffected parents and not transmitted to child with OFC are shown below. Variants tested in zebrafish assays are bolded. (**B**) Levels of *krt4* expression as a read out for functional validation of three human *ZNF750* variants in a zebrafish assay. Zebrafish embryos (5.5 hpf) were injected individually with control (Ctrl) or ZNF750 morpholinos (MO), then injected with: LacZ; human RNA encoding *ZNF750* variants (p.Ser160Pro, p.Cys164Ser, p.Pro646Leu); or the human reference sequence (Ref). Significant differences in expression are denoted following One-way ANOVA; \**P* < 0.05;\*\**P* < 0.01. Only significant differences are shown. Data are means of 4 biological replicates +standard deviations.

Because functional predictions do not always match results from embryo-based functional tests (Li et al. 2017), we used a zebrafish-embryo-based assay to test the function of transmitted *ZNF750* variants that are predicted to be damaging (p.Ser160Pro and p.Pro646Leu) and benign (p.Cys464Ser). RNA-sequencing (RNA-seq) of *znf750* morpholino oligonucleotide (MO)-injected fish embryos demonstrate effective knockdown of wild-type *znf750*, and that expression of *krt4*, a periderm marker in zebrafish, is among the most down-regulated relative to control-MO injected embryos (Appendix Table 2). Of note, *znf750* MO-injected embryos, observed up until 24 hpf, do not exhibit a gross phenotype. To test the function of the three variants, we first injected zebrafish embryos with *znf750* MO and then injected them with mRNA encoding variants of human *ZNF750* (reference, p.Ser160Pro, p.Cys464Ser, or p.Pro646Leu), mouse *Zfp750* (reference, and the orthologous residue p.Ser154Pro), or encoding Beta galactosidase (*LacZ*) as a control (Fig. 1B, Appendix Fig. 1). Embryos injected with *znf750* MO and the negative-control *lacZ* mRNA showed a 50% reduction of *krt4* expression relative to embryos injected with the control MO, as expected from the RNA-seq results (Fig. 1B). By contrast, mRNA encoding the *ZNF75*0 reference variant more than reversed the effects of the MO (Fig.1B). All of the patient-derived *ZNF750* variants were able to elevate *krt4* expression more effectively than the negative control protein, but the two variants predicted to be damaging were significantly less effective, and near significantly less effective (*P* = 0.0503), respectively, than the reference variant (Fig. 1B). These results concord with the algorithms’ predictions.

As a further test, we engineered murine knock-in alleles of the orthologous residues, i.e. *Zfp750* p.Ser154Pro (*Zfp750*^*S154P*^) and p.Pro626Leu (*Zfp750*^*P626L*^). We found that mosaic F0 crispants composed largely of cells homozygous for *Zfp750*^*S154P*^ or *Zfp750*^*P626L*^ developed normally. For *Zfp750*^*S154P*^ we additionally generated a stable line, which is not mosaic. In the F2 and subsequent generations, *Zfp750*^*S154P/S154P*^ embryos also developed normally. In addition, oral adhesions were absent from such embryos at E15.5 (Appendix Fig. 2A, B). We then conducted RNA-seq on forelimbs from *Zfp750*^*S154P/S154P*^ embryos and compared their gene expression levels with the ones from wild-type at E15.5. Very few genes were differentially expressed in *Zfp750*^*S154P/S154P*^ relative to in wildtype forelimbs, and such genes were not enriched for any gene ontology (GO) terms (Appendix Table 3). Consistent with the absence of a phenotype in *Zfp750*^*S154P/S154P*^, mRNAs encoding the wild-type or p.Ser154Pro variants of murine Zfp750 were equally effective at elevating *krt4* expression in zebrafish embryos depleted of *znf750* (Appendix Fig. 1); these results contrast with those using variants of human *ZNF750*. In summary, murine embryos homozygous for variants orthologous to p.Ser160Pro and p.Pro646Leu did not exhibit gross phenotypes.

### Zfp750 is necessary for epidermal differentiation, temporary fusions, permeability barrier and embryonic survival

To fully examine a putative role for ZNF750 during orofacial development, we used CRISPR/Cas9 editing to introduce a frame shift indel into the *Zfp750* gene (Fig. 2A). *Znf750*^*fs/fs*^ were detected in Mendelian ratios (Appendix Fig. 3). However, pups died in the early perinatal period, perhaps due to dehydration. At E18.5, *Znf750*^*fs/fs*^ embryos present macroscopic defects compared to wild-type littermates (Fig. 2B, C) including open eyes (Fig. 2D, E), unfused pinnae (Fig. 2F, G), and unfused digits (Fig. 2H, I). Mutant embryos also show increased permeability to toluidine blue relative to wildtypes (Fig. 2J, K) indicating an impaired epidermal barrier, as reported earlier (Butera et al. 2023; Schwartz et al. 2024).

**Figure 2:**
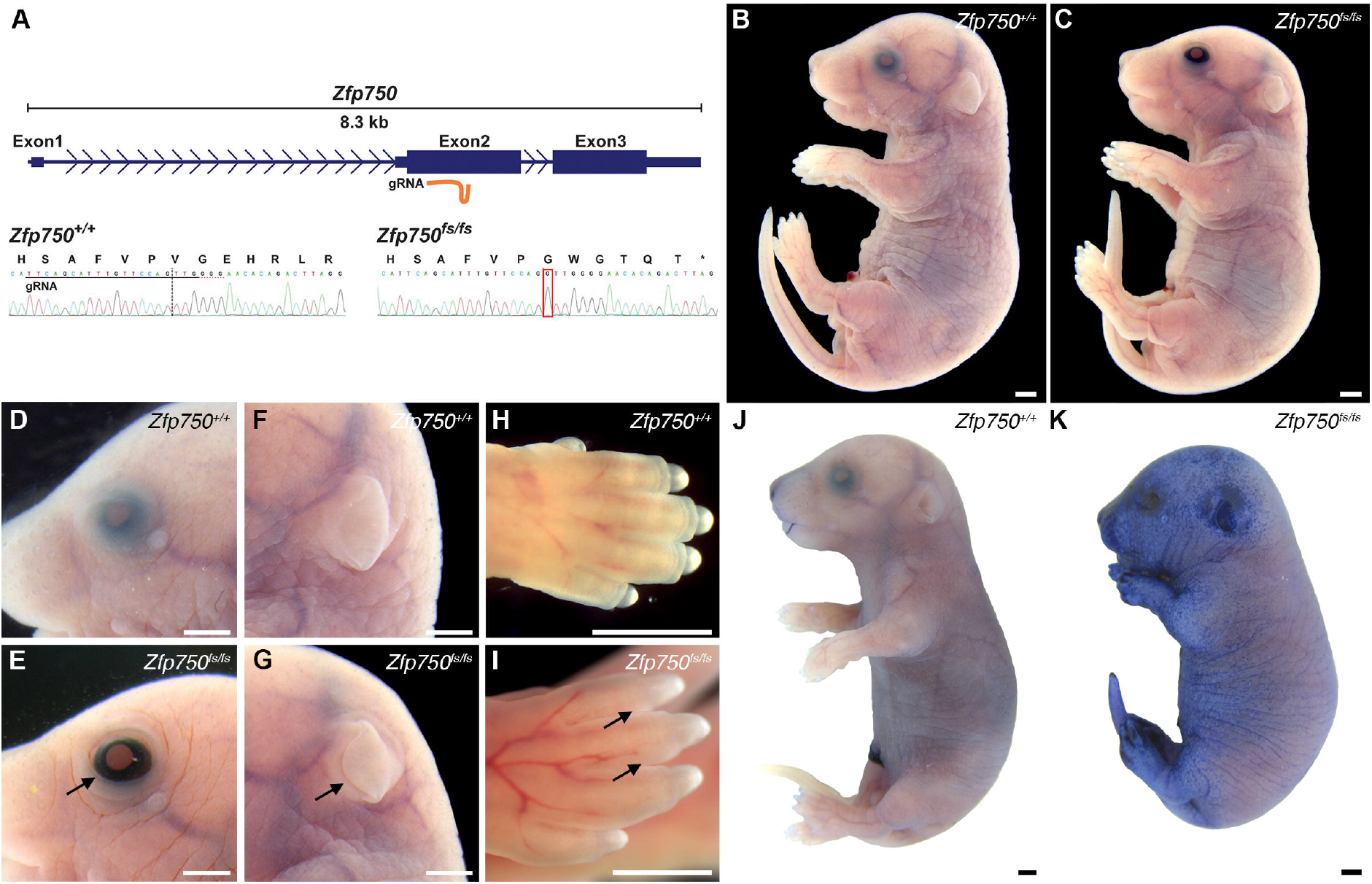
Frameshift mutation in *Zfp750* leads to periderm-associated embryonic defects and lack of epidermal barrier function. (**A**) Schematic of *Zfp750* allele and frameshift mutation in Exon 2. Representative Sanger sequencing data from wild-type and *Zfp750*^*fs/fs*^ embryos shows the +1 G insertion (red box) and subsequent premature STOP codon (asterisk) created in mutants. Position of the Guide (g) RNA is indicated below Exon2 (top panel) and below the wild-type sequence. (**B-I**) Macroscopic views of wild-type (B, D, F, H) and *Znf750*^*fs/fs*^ embryos (C, E, G, I) at E18.5. Black arrows point to open eyes, unfused pinnae and unfused digits in mutants. (E, G, I) Dye exclusion assay of whole embryos show a functional barrier in wild-type (J) but not in *Zfp750*^*fs/fs*^ (K) embryos. Scale bars: 1 mm

### Absence of Zfp750 alters the epidermal gene expression profile of embryonic forelimbs

To identify differentially expressed genes that might explain the gross phenotypes of *Zfp750*^*fs/fs*^ embryos, we conducted RNA-seq on right paws from E15.5 embryonic forelimbs (Fig. 3). We found a total of 568 genes significantly differentially regulated (absolute log2-fold change expression > 0.5 and *P*-adj < 0.05), with 62 being up-regulated and 506 being down-regulated in *Zfp750*^*fs/fs*^ tissues relative to widltypes (Fig. 3A, Appendix Table 4). Up-regulated genes are enriched for GO terms related to epidermal development, whereas down-regulated genes are enriched for terms related to epidermal differentiation, with the strongest effect size on genes associated with the cornified envelope (Fig. 3B, C, Appendix Figs. 4, 5). Up-regulated genes include *Tp63*, a marker of the basal layer, and *Cldn7* and *Cldn8*, components of tight junctions (Fig. 3B, C, Appendix Table 4). Down-regulated genes include many involved in terminal epidermal differentiation (e.g., *Lor, Sprr3, Flg2*), lipid synthesis (e.g., *Alox12b, Aloxe3*)(Fig. 3C), and desmosome structure (e.g., *Dsc1*). Among genes whose expression is relatively specific to periderm cells compared to other epidermal cells at E14.5 (Jacob et al. 2023), three are down-regulated at least two-fold (*Krt19, Paqr6, Cldn23*) and two are up-regulated (*Krt8, Krt17*) in *Zfp750*^*fs/fs*^ versus in wild-type embryonic paws (Fig. 3C,D). We conclude that Zfp750 promotes terminal epidermal differentiation and is required for proper gene expression in the periderm.

**Figure 3:**
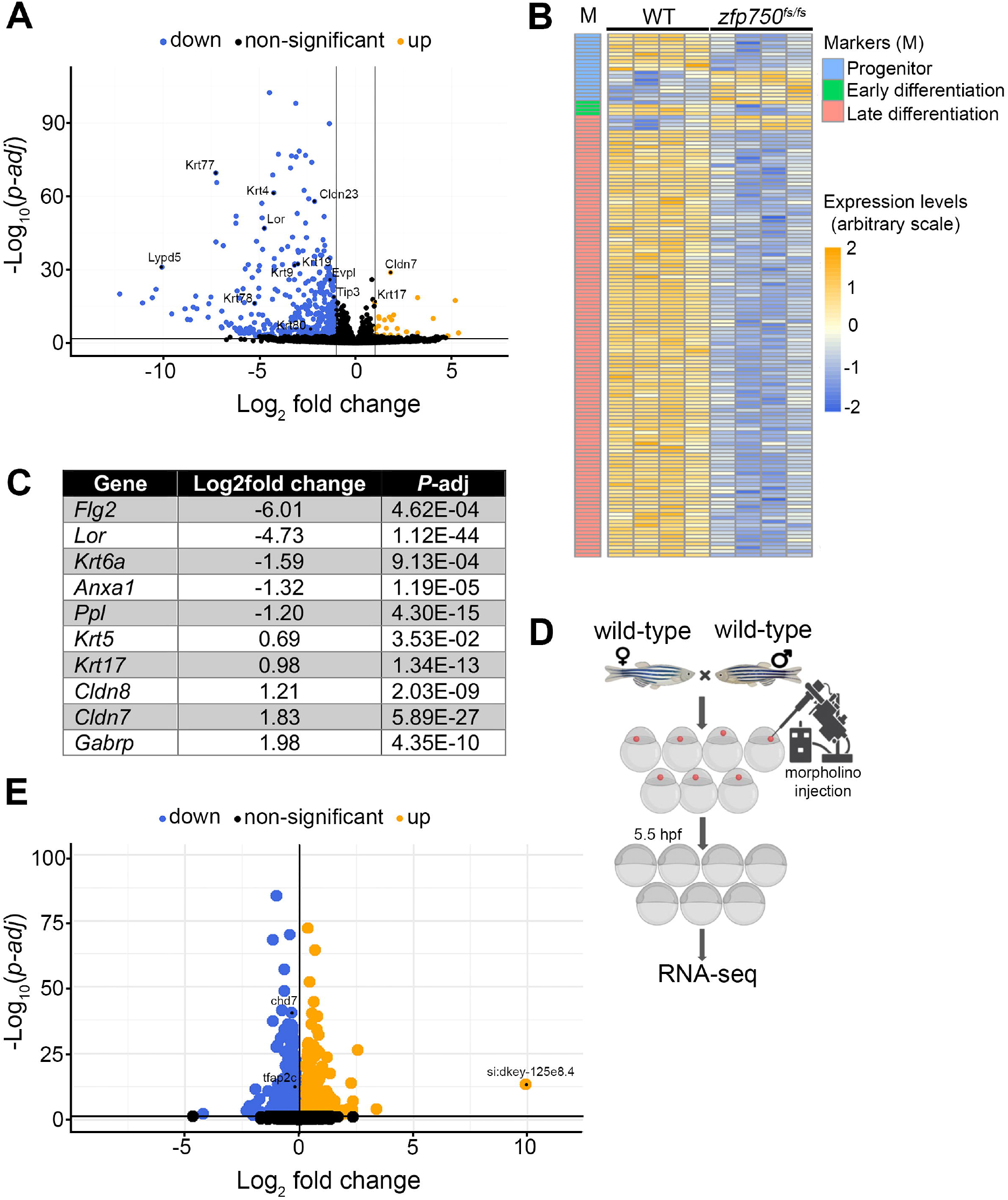
Disruption of *Zfp750 (*and *znf750*) alters differentiation and periderm-related genes in mouse and zebrafish embryos. (**A**) Volcano plot showing log2 fold change (x-axis) in gene expression between *Zfp750*^*fs/fs*^ murine paw and corresponding wild-type control at E15.5. Adjusted *P*-values from the comparisons are plotted on y-axis. Upregulated, downregulated, and non-significantly changed gene expression are shown as orange, blue, and black dots, respectively. (**B**) Heatmap of normalized (and scaled) expression level of differentiation-related genes (early, late, and progenitor) that are differentially expressed between *Zfp750*^*fs/fs*^ murine paw and corresponding wild-type control at E15.5. (**C**) Table showing genes whose expression is enriched in periderm relative to in other tissues and that are differentially expressed in *Zfp750*^*fs/fs*^ murine paws compared to in wild-type paws. (**D**) Schematic of *znf750* morpholino injections in zebrafish embryos and sample collection for RNA-sequencing. (**E**) Volcano plot showing log2 fold change (x-axis) in gene expression between *znf750* morpholino (MO) injected zebrafish embryos and standard control MO injected control embryos at 5.5 hpf. Adjusted *P*-values from the comparisons are plotted on y-axis. Genes with up, down, and non-significant gene expression are shown as orange, blue, and black dots respectively.

### Interdigital ridges are absent in Zfp750^fs/fs^ embryos

The separation of the digits, which occurs between E13.5 to E15.5, depends of migration of epidermis into a zone of dying mesenchymal cells (Kashgari et al. 2020). Around E16, immediately after the digits have separated, they begin to fuse to one another in a proximal to distal progression and remain fused until a few days after birth (Maconnachie 1979). A transient fusion resolving after birth also occurs between eyelids and between pinnae and scalp (Maconnachie 1979); the digits, eyelids and pinnae are all examples of *temporary epithelial fusions* (Mazzalupo and Coulombe 2001). Interdigital ridges are present in wild-type (Fig. 4A,B) but not in mutant forelimbs (Fig. 4C,D) and hindlimbs (Appendix Fig. 6). Because periderm cells are proposed to contribute interdigital ridges (Maconnachie 1979), we examined the morphology of periderm cells by scanning electron microscopy. On the surface of digits in wild-type embryos periderm cells have a cobblestone appearance, with obvious protruding nuclei, and microvilli on their apical surface (Fig. 4B,E). In *Zfp750*^*fs/fs*^ embryos, periderm cells have less prominent nuclei and fewer microvilli (Fig. 4D,F). We performed immunofluorescence microscopy on transverse sections of forelimb digits containing interdigital ridges in wild-type embryos (Fig. 4E-J) and through the equivalent position along the proximal-distal axis of digits in *Zfp750*^*fs/fs*^ embryos, which lack these structures. E-cadherin, a marker of adherens junctions, is present in epithelial cells covering the digits (Fig. 4E, F, Appendix Fig. 7) and Keratin (K) 14 in all basal cells and occasional suprabasal cells but not in the most superficial cells in both genotypes (Fig. 4G, H). K14 expression is also prominent in suprabasal cells of the interdigital ridge in wildtypes (Fig.4G). Periderm markers, including K6 (Mazzalupo and Coulombe 2001), K17 (McGowan and Coulombe 1998), K8/18 (Lu et al. 2005), and Annexin1 (Fava et al. 1993), are expressed in the most superficial epithelial cells covering the digits in both genotypes (Fig. 4E-H’, Appendix Fig. 7), and in epithelial cells of the interdigital ridge in wild types only (Fig. 4E, E’, G, G’, Appendix Fig. 7). While K18 is expressed in all layers of the epidermis in both genotypes, it appears more diffuse and less intense in wildtypes than in mutants (Fig. 4J, L). Lastly, Loricrin, a protein characteristic of terminal epidermal differentiation, is detected in suprabasal cells and in cells overlying (dorsally) the fused wild-type embryonic digits (Fig. 4I) but not in *Zfp750*^*fs/fs*^ embryos (Fig. 4J) as previously reported (Butera et al. 2023; Schwartz et al. 2024). We conclude that *Zfp750* is required for terminal epidermal differentiation, for normal gene expression in the periderm, and for the formation of interdigital ridges.

**Figure 4:**
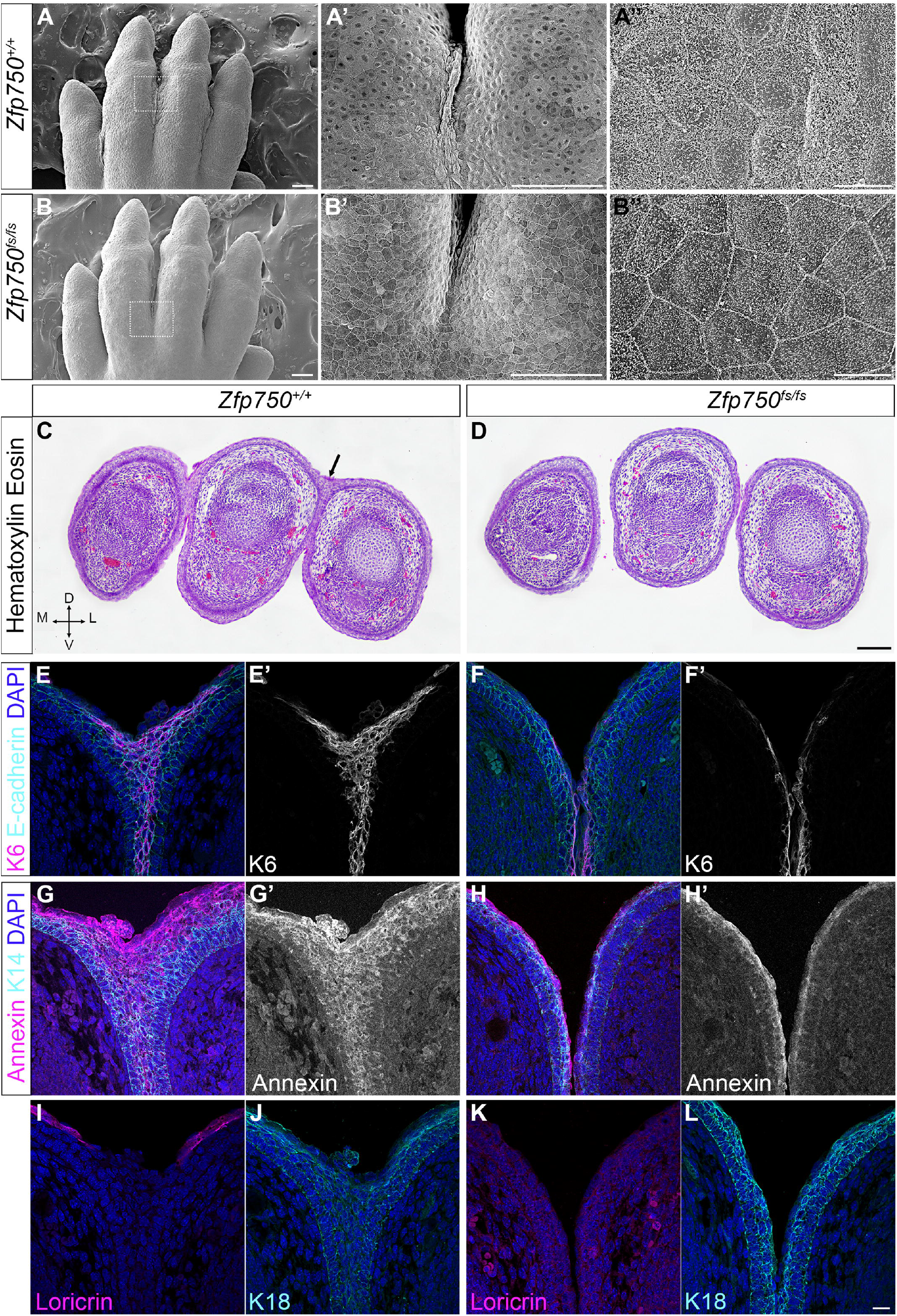
Frameshift mutation in *Zfp750* leads to absence of interdigital ridge despite the presence of periderm-associated proteins. (**A-B’’**) Scanning electron microscopy and (**C, D**) hematoxylin and eosin-stained transverse sections of E15.5 forelimbs reveal the presence of interdigital ridge (A [zoom-in of white boxes in A’], black arrow in C) and epidermal microvilli on the surface of wild-type (A’’) but not in *Zfp750*^*fs/fs*^ embryos (B, [zoom-in of white boxes in B’], B’’). (**E-L**) Immunofluorescent staining of wildtype (left two columns) and *Zfp750*^*fs/fs*^ (right two columns) for E-cadherin and Keratin 6 (dual staining in color, single K6 channel shown in black and white), Keratin 14 and Annexin A1 (dual staining in color, single Annexin channel shown in black and white), Loricrin, and K18. Proteins against which antibodies were directed are indicated. Nuclei are counterstained with Hoechst (blue). Overall region stained is shown in black boxes in C and D. Scale bars: A-B’, C, D = 100 µm; A”, B” = 10 µm; E-L = 20 µm. D = dorsal, V = ventral, M = medial, L = lateral.

### Znf750 is downstream of Grhl3 in the zebrafish enveloping layer transcriptional regulatory network

The altered peridermal phenotype in *Zfp750*^*fs/fs*^ may reflect periderm-autonomous function of *Zfp750* or an indirect effect of *Zfp750* function in ectodermal cells, from which the periderm is derived (Holbrook and Odland 1975). To determine if ZNF750 is involved in gene regulation within periderm cells, we examined its role in the zebrafish enveloping layer, the periderm-equivalent that derives from early blastomeres rather than from ectoderm (Kimmel and Law 1985). We injected zebrafish embryos at the 1-2 cell stage with *grhl3* MO, or a standard control MO, harvested RNA from embryos at 5.5 hpf and conducted RNA-seq. Expression of *znf750* is strongly down regulated in *grhl3* morphants relative to in control MO-injected embryos. By contrast, *grhl3* expression is unchanged in *znf750* morphants relative to control MO-injected embryos (Appendix File 1). Importantly, there are many more differentially expressed genes in *grhl3* morphants than in *znf750* morphants (Appendix File 1). These results place *Znf750* downstream of *Grhl3* in the fish periderm transcriptional regulatory network.

### Palatal shelves fuse but oral adhesions are present in Zfp750^fs/fs^ embryos

Histological analysis of serial coronal sections through the head of wild-type and *Zfp750*^*fs/fs*^ embryos at E15.5 reveals fused palatal shelves and the absence of epithelial cells in the midline in all wild-type and *Zfp750*^*fs/fs*^ embryos (Fig. 5A-D’). E-cadherin and K6 (Fig. 5C-D’), Annexin 1 and K14 (Fig. 5E-F’), K17 (Appendix Fig. 7), and K18 (Fig. 5H, J) are detected in the oral epithelium and periderm in the same pattern as in the forelimb epidermis. Loricrin is absent from both wildtypes and *Zfp750*^*fs/fs*^ embryos (Fig. 5G, I) because terminal differentiation is not yet initiated at this developmental stage in the oral cavity. In both wild types and *Zfp750*^*fs/fs*^, epithelial triangles are visible on the oral side of the palatal shelf fusion zone, reflecting cells that have migrated from the midline (Teng et al. 2022). Therefore, Zfp750 is dispensable for the behavior of medial-edge epithelial cells, including periderm cells, during palatal fusion in murine embryos.

**Figure 5:**
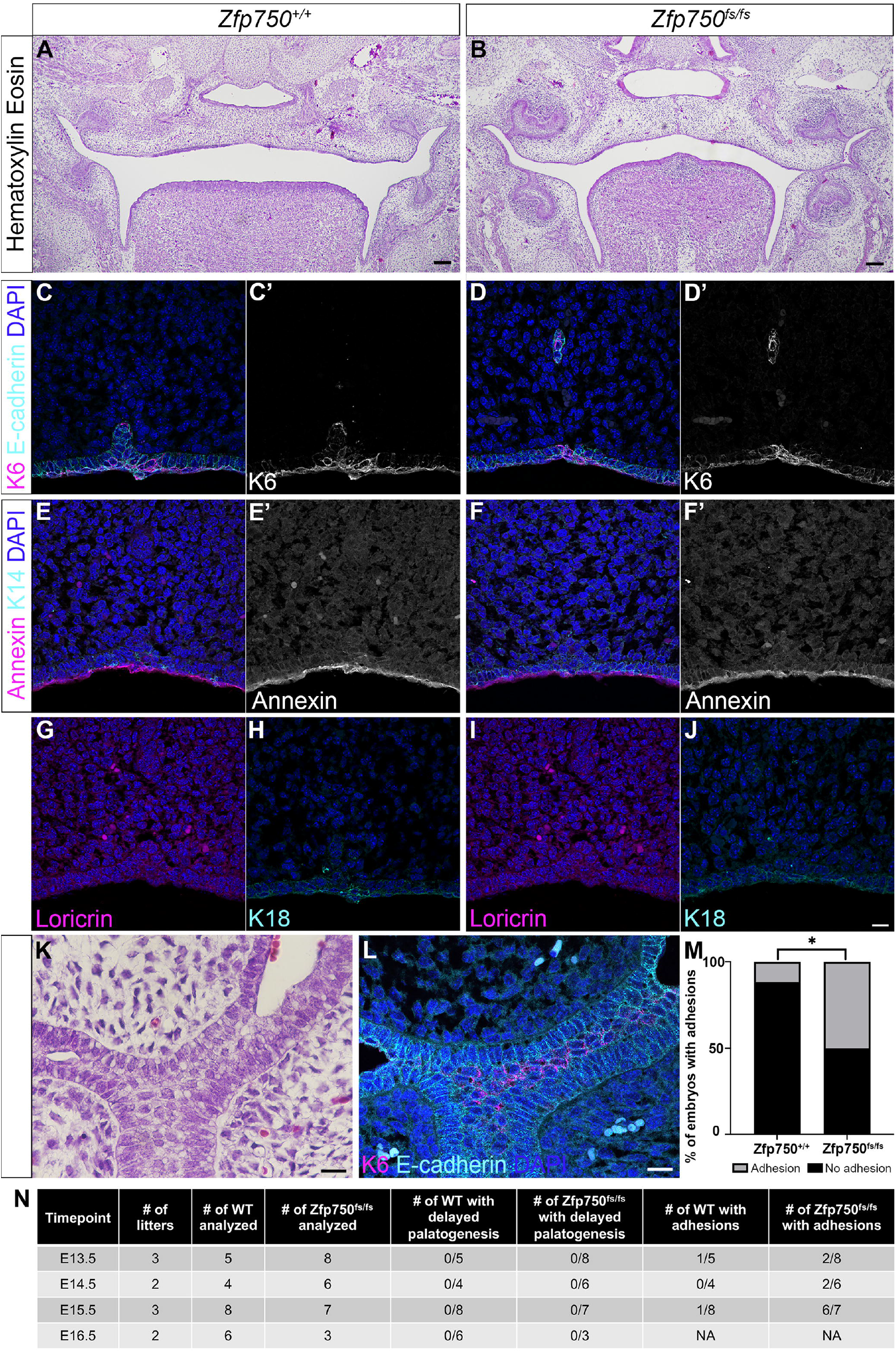
Oral adhesions, but properly fused palatal shelves, are present in *Zfp750*^*fs/fs*^ mutant embryos. (**A-B**) Hematoxylin and eosin stained coronal sections of wild-type (A) and *Zfp750*^*fs/fs*^ E14.5 embryos (B). (**C-N**) E14.5 coronal wildtype (left two columns) and *Zfp750*^*fs/fs*^ (right two columns) sections immunofluorecently stained for E-cadherin and Keratin 6 (dual staining in color, single K6 channel shown in black and white), Keratin 14 and Annexin A1 (dual staining in color, single Annexin channel shown in black and white), Loricrin, and K18. Proteins against which antibodies were directed are indicated. Nuclei are counterstained with Hoechst (blue). (K) Hematoxylin and eosin stained coronal section of a E14.5 *Zfp750*^*fs/fs*^ embryo with intra-oral tissue-tissue adhesion between the tooth bud and the maxilla. Adjacent section dual stained for Keratin 6 and E-cadherin is shown in L. (M) Quantification of the percentage of embryos that display tissue adhesions. (N) Table displaying the incidence of oral adhesions observed in wild-type and *Zfp750*^*fs/fs*^ embryos at several embryonic developmental time points. Scale bars: A, B = 100 µm; C-L = 20 µm.

In several murine models of OFC genes, palatal shelves fuses but there is ununusally high incidence of oral adhesions (Paul et al. 2017; Peyrard-Janvid et al. 2014). In *Zfp750*^*fs/fs*^ embryos we observe tissue adhesions between the oral epithelium of the maxilla and of the mandible (Fig. 5K), or between the oral epithelium of the maxilla and the lingual epithelium, at a higher frequency than in wildtypes (Fig. 5M, N). We observed more oral adhesions in *Zfp750*^*fs/fs*^ embryos at E16.5 than at E13.5. The periderm-specific protein K6 is detected within the sites of adhesion (Fig. 5L) as reported in *Irf6* heterozygous mutants (Kousa et al. 2017). We conclude that loss of Zfp750 disrupts the anti-adhesive quality of the periderm, at least within the oral cavity.

## Discussion

Here we tested the hypothesis that *ZNF750* is an OFC risk gene. The rationale for this hypothesis can be summarized as “guilt by association” as *ZNF750* shares some properties with other genes in its regulatory network that are also involved in OFCs. We assessed whether *ZNF750* variants could be found in families with VWS who have no detectable mutation in *IRF6, GRHL3*, or *PRKCI* but did not find any variants in those families. We also sequenced 2,125 individuals with nsOFCs and their family members. In contrast to the types of variants found for *IRF6, GRHL3*, and other genes (*de novo* and truncating variants), we only found rare missense variants, which are more difficult to interpret. We tested two of these variants that were predicted to be damaging, and found that both had lower activity than the reference variant in a functional zebrafish embryo-based assay. However, murine embryos homozygous for orthologous variants did not exhibit the gross phenotypes. Together, these studies indicate that damaging coding variants in *ZNF750* are not frequently detected in nsOFC patients. In contrast, there is strong genetic evidence that variants in *ZNF750* are associated with a dominantly inherited dermatosis phenotype (Birnbaum et al. 2011). We are not aware of cutaneous phenotypes in the individuals from our cohorts, but onset of these phenotypes is often well after the age of recruitment for OFC. Similarly, there is no mention of OFCs in the published *ZNF750* families. It remains possible that certain moderate-effect alleles of *ZNF750* modify risk for OFCs, which would require more systematic testing of missense variants. In an era where rare diseases are increasingly being recognized and diagnosed molecularly, it is also possible that a rare syndrome characterized primarily by cutaneous lesions but with incompletely penetrant craniofacial phenotypes exists and is caused by *ZNF750* mutations.

We examined murine embryos homozygous for a frame-shift in *Zfp750* for cleft palate and oral adhesions. The presence of oral adhesions together with a fused palate have become hallmarks of murine models of OFC-risk genes (Ingraham et al. 2006; Kousa et al. 2017; Paul et al. 2017; Peyrard-Janvid et al. 2014; Richardson et al. 2009). Expression of proteins characteristic of periderm within the oral adhesions of *Zfp750* mutants is also seen in murine models of OFC genes (heterozygotes for a loss-of-function (LOF) allele of *Irf6* (Kousa et al. 2017), *Arhgap29*^*K326X*^ (Paul et al. 2017), and *Grhl3*^*LOF*^ (Peyrard-Janvid et al. 2014). However, we found that 0 of 10 *Zfp750*^*fs/fs*^ embryos at E15.5 exbhibited a cleft palate, whereas 1 of 6 *Grhl3*^*LOF/LOF*^ did (Peyrard-Janvid et al. 2014). The *Zfp750* phenotype is less severe than that of *Grhl3*^*LOF/LOF*^ because while oral adhesions are present in both genotypes, syndactyly only occurs in the latter. This is consistent with the observation that *Znf750* appears to function downstream of *Grhl3*. Of note, milder phenotypes for mutants for genes lower in the regulatory network is also seen in human peridermopathies. Genes higher in the network are associated with more severe syndromes that include OFC, but also include structural abnormalities in other organs. In contrast, individuals with variants in *GRHL3* diagnosed with VWS, have a less penetrant phenotype and are less likely to have lip pits, digit, and dental anomalies than those with *IRF6* mutations (Peyrard-Janvid et al. 2014). In conclusion, although the shared phenotype of oral adhesions in *Zfp750* and *Grhl3* mutants lends some support to the hypothesis that *ZNF750* variants, like *GRHL3* variants, can contribute risk to nsOFC, the milder phenotype of *Zfp750* mutants lessens the certainty of this conclusion.

Finally, the presence of oral adhesions but the absence of temporary epithelial fusions in *Zfp750* mutants adds complexity to our understanding of the role of the periderm in embryonic fusion and septation events. The periderm is accepted to have anti-adhesive function that prevents unwanted fusion between embryonic structures of the oral cavity and aids in the septation of digits (Kashgari et al. 2020). Nonetheless, temporary epithelial fusions may be facilitated by the periderm. Clusters of periderm cells are present adjacent to transiently fused digits and eyelids; it has been proposed these clusters result from periderm migration (Maconnachie 1979). However our finding of periderm markers between transiently fused digits in wild types implies it is not necessary for periderm to completely evacuate the space between digits for them to temporarily fuse. Instead, the absence of temporary epithelial fusions in *Zfp750*^*fs/fs*^ supports the possibility that Zfp750 is required for a pro-adhesive function of the periderm. Future experiments are necessary to distinguish pro-adhesvie and anti-adhesive functions of periderm in distinct developmental and anatomical contexts.

## Acknowledgements and funding

This works was funded by grants from the National Insitute of Health (R01DE028300 to AB; R01DE027983 (PI:Eric Liao) and DE023575 to RAC; F31DE033222 and T32GM145441 to EA; R03TR004674 to MD; U54OD030187 and R01DE032319 to SAM). We are grateful for technical support from Gregory Bonde, Frank Radella, Josh Rosswork, Jiarui Jiang, Abby Kay, and Johnny Jianbao. We acknowledge the support from the Central Microscopy Research Facility at the University of Iowa for scanning electron microscopy. The Jackson Laboratory Genetic Engineering Technology core is supported in part by the JAX Cancer Center (P30CA034196).

## Author contributions

S. Singh, E. Adelizzi, M., A. Butali, E.J. Leslie-Clarkson, M. Dunnwald, and R.A. Cornell contributed to conception and design of the study, data acquisition, analysis, and interpretation. S. Singh, E. Adelizzi, M., E.J. Leslie-Clarkson, M. Dunnwald, and R.A. Cornell drafted and revised the manuscript; C. Heffner, S.A. Murray, contributed to data acquisition and analysis; K. Duncan, S. Curtis, W. Awotoye, J. Olotu, T. Busch, W.L. Adeyemo, J.J.L. Gowans, T. Naicker contributed to data acquisition and analysis. All authors gave final approval and agree to be accountable for all aspects of the work.

## Data availability statement

Data presented in this work are available at NCBI SRA SUB15933886. All authors gave final approval and agree to be accountable for all aspects of the work.

## Methods

The detailed methodology is described in the Appendix Materials and Methods.

