## Appendix for "Zfp750 prevents oral adhesions and promotes temporary epithelial fusions"

**Material and Methods**

*Sequencing of cohorts with OFCs*

Whole-genome sequencing from 5,509 individuals from the Gabriella Miller Kids First Program was analyzed for variants in *ZNF750* described in (Bishop et al. 2020; Curtis et al. 2025; Mukhopadhyay et al. 2020). This dataset included 1,851 affected probands and their parents. Variants in the *ZNF750* region were annotated with VEP (McLaren et al. 2016). The sequencing was conducted by the Broad Institute, and data were aligned to hg38 and called with GATK pipelines (McKenna et al. 2010; Van der Auwera et al. 2013). Quality control for individuals included removing samples with a missingness value, Mendelian error value, or transition/transversion (Ts/Tv) ratio outside of three standard deviations from the mean. Family relationships and sex were confirmed by calculating identity-by-descent and X-chromosome heterozygosity with PLINK (v1.90) (Chang et al. 2015). Biallelic variants were then filtered for minor allele count (MAC) ≥ 1, GQ ≥ 20, DP ≥ 10, missingness $\leq$ 5% using VCFTools (v0.1.13). Variants with a quality by depth (QD) $\leq$4 or that deviated from Hardy-Weinberg equilibrium (*P* < 1$\times$10^-7^) were excluded. Variants were annotated for minor allele frequence and consequence using VEP (McLaren et al. 2016).

We used direct sequencing to search for sequence variations in coding regions of the *ZNF750* (NM_024702 3 exons) in 150 African individuals with CLP. Primers were designed using Primer 3 and optimized to the optimal annealing temperature. Details available on request through the Butali lab website (https://butali.lab.uiowa.edu/). We sent PCR products for sequencing using an ABI 3730XL (Functional Biosciences, Inc., Madison, WI), transferred Chromatograms to a Unix workstation, base-called with PHRED (v.0.961028), assembled with PHRAP (v. 0.960731), scanned by POLYPHRED (v. 0.970312). All sequence results were viewed with the CONSED program (v. 4.0). When available, we sequenced samples from the mother and father to confirm if variants are *de novo* or segregating in the family. We then compared the variants found in this study to variants that are present in the African population in gnomAD version 4.1 (Karczewski et al. 2020). The functional effects of these variants on the protein were predicted using polyphen (<http://genetics.bwh.harvar-d.edu/pph2/>) (Adzhubei et al. 2013), and SIFT (<http://sift.jcvi.org/>) (Ng and Henikoff 2003).

Variants were considered rare if they had a minor allele frequency less than 0.1% in any gnomAD population. Variants not found in gnomAD were considered novel. Variants were considered possibly damaging if they were predicted to be damaging by either Polyphen or SIFT.

*TDT Burden Tests*

A burden of rare variants in *ZNF750* region was tested using a rare variant extension of the transmission disequilibrium test (RV-TDT) (He et al. 2014). We analyzed data from a subset of 759 complete trios with OFCs from the Gabriella Miller dataset described above and included all rare variants with MAF < 0.1%. *P* < 0.05 was considered statistically significant.

*Animal use*

All the procedures carried out in this project involving zebrafish were approved by the Institutional Animal Care and Use Committee (IACUC) of the University of Iowa and the University of Washington. All mouse procedures were conducted according to national and international guidelines (AALAC and IACUC) and have been approved by the Jackson Laboratory Animal Care and Use Committee.

*Zebrafish embryo micro-injections*

*ZNF750* (and *Zfp750*) reference and variants were synthesized and cloned in pCS2^+^ vector by GenScript (<https://www.genscript.com/>). *ZNF750* (and *Zfp750*) variants were generated by 1-bp site-directed mutagenesis in the reference gene sequence. Reference and variant-containing mRNAs were synthesized from 1 μg of linearized plasmids by using the mMESSAGE mMACHINE SP6 Transcription kit (Invitrogen; cat. #AM1340). mRNA products were treated with DNase (Invitrogen DNA-free DNA Removal kit; cat. #AM1906) and purified using RNA Clean & Concentrator-25 kit (Zymo Research; cat. #R1017). Normal looking zebrafish embryos that had undergone at least one but fewer than four cell divisions (i.e., 2-8 cell stage) were injected with approximately 0.1 ng of splice-blocking morpholino against zebrafish *znf750* (5’-ACTTAGATTTTACCTACCTTTCTGT-3’; GeneTools Inc.)*,* standard control morpholino (5'-CCTCTTACCTCAGTTACAATTTATA-3'), and/or approximately 100 pg mRNAs encoding *Escherichia coli* β-Galactosidase (*lacZ*), or human *ZNF750* reference, or individual human patient variants, or mouse *Zfp750* reference, or *Zfp750* p.Ser154Pro variant. Injected embryos were kept in embryo medium (5.03 mM NaCl, 0.17 mM KCl, 0.33 mM CaCl_2_, 0.33 mM MgSO_4_, 0.1% Methylene blue) at 28.5°C until harvested for RNA extractions.

For qRT-PCR wild-type embryos from the AB strain were used. For RNA seq, wild-type outbred embryos from commercially-obtained fish were used. Approximately 40 morpholino- and mRNA-injected 5.5 hpf embryos per group were used for total RNA extraction using RNAqueous Total RNA Isolation kit (Invitrogen, cat. #AM1912). All living embryos, regardless of morphology, were included in the analysis of mRNA levels. For cDNA preparation, 1 μg of DNase-treated total RNA was processed with High-Capacity cDNA Reverse Transcription kit (Applied Biosystems, cat. #4368814). Real-Time PCR assays were performed to measure zebrafish keratin 4 (*krt4)* and glucose-6-phosphate dehydrogenase (*g6pd*) genes expression by using 10-fold-diluted cDNA samples and iTaq Universal SYBR Green Supermix (Bio-Rad, cat. #1725121) on the CFX Connect Real-Time PCR Detection System (Bio-Rad; cat. #1855201). Zebrafish actin gene (*actb1)* was used as reference gene to calculate the fold change in *krt4* gene expression by using the 2−ΔΔCt comparative Ct method (Livak and Schmittgen 2001). It is potentially a cofounder that we invariably injected *znf750* morpholino before we injected mRNAs, but this seems unlikely. To avoid a confounder of the order in which the mRNAs were injected, in the four replicates of the experiment, we injected the mRNAs (i.e., negative control lacZ mRNA, reference variant of *ZNF750*, patient variants of *ZNF750*) in different orders. The person performing qRT-PCR was not blinded to the experimental groups they evaluated. Primers used are as follow:

| **Primer** | **Sequence (5' - 3"** |
| --- | --- |
| RT-qPCR krt4 Forward | GACAACAGCAGAAACCTGGA |
| RT-qPCR krt4 Reverse | GCAATCTCAGCCTTTGTTGA |
| RT-qPCR g6pd Forward | TGCGGGATGAGAAGGTAAAAGT |
| RT-qPCR g6pd Reverse | GATCTCCGACATACTGACCCAG |

The relative expression levels of *krt4* are presented as the average of four biological replicates of at least 40 embryos each. Using four biological replicates instead of three increases the sensitivity to detect small differences in gene expression between groups. One-way ANOVA with Dunnett correction for multiple comparisons was performed between treatment groups. ANOVA and related tests are parametric which assume data follow a specific distribution, with known parameters (mean, standard deviation). Because of the small sample size, we were unable to confirm that the distribution of data was normal, but there is no a priori reason to assume otherwise.

*Generation of mutant alleles in F0 mouse embryos*

Candidate guide RNA (gRNA) sequences for CRISPR/Cas9 editing were selected based on efficiency and off-target scores, as well as proximity to the orthologous human variant SNP. Based on these criteria, guides *Zfp750*-Ser154Pro-FWD 5’-TTCAGCATTTGTTCCAGTTG-3’ (forward strand) and *Zfp750*-Pro626Leu-FWD (forward strand) 5’-CAGCCCAGGGAACGTCCGAG-3’ were selected as the gRNA sequences for the Ser154 and Pro626 mutations, respectively. A 200-bp single-stranded DNA (ssDNA) donor oligonucleotide was designed for each variant. ssDNA donors contained both the SNP variant as well as a silent mutation in the PAM sequence to prevent re-cutting after HDR events. ssDNA Donor Sequences are as follow:

p.Ser154Pro variant: [5’ - GGAATGAGGAGGCTTTGGTGGTAGAGTTGGCCAGCACTTCAGTGGCCTCTGTATCTTCTGGCCCCCTAAGTCTGTGTTC**A**CCAACTGGAACAAATGCTG**G**ATGTCTAACCCCACTGTCGAGGCAGGGCTGGGTGCTGAGAATGGCTTCTGGTGCCATTTCTTTCTGTAGGGCTGGTTTCTGGGGACCTTTGTGGGCTCCC – 3’ (reverse strand)]

p.Pro626Leu variant: [5’ - CACTGAGCCTCCAGGTTCTCTGTAGCACTGAGGGTGGGCACATCCTGACGGGTGGGTTCCTGGACTGTGCCTTCCTCATCGGC**G**ACTCGGACGTTCCCT**A**GGCTGTAGGCCGCCAGCTGACAGAGAGCTACAGCTGCCGTCTGCTTCTGTTCATCACTGCTGTCCTCCGTGTGCACGTCTGGGGTTGTCCTGCTGTGGGC– 3’ (reverse strand)]

The crRNA, trRNA and ssDNA donor oligos were sourced from Integrated DNA Technologies (IDT). Zfp750-Ser154Pro and Zfp750-Pro626Leu crRNAs were annealed with trRNA following IDT Alt-R System protocols. Briefly, both components were resuspended at 100 μM in IDT Duplex Buffer, combined in equal amounts, heated to 95°C for 5 min, and allowed to cool passively to room temperature. Following this annealing step, the concentration of annealed guide RNA was assayed by NanoDrop. The Crispr:tracr guide RNA hybrid was complexed with Alt-R Cas9 at 37°C for 15 min in a thermocycler.

For electroporation, zygotes from C57BL/6NJ mice raised at the Jackson Labs were harvested and placed in a droplet comprised of 10 μl TE buffer, 10 μl Opti-MEM (Millipore Sigma), ssDNA donor (2000 ng/μl) and the ribonucleoprotein complex containing Alt-R Cas9 nuclease (500 ng/μl) and gRNA (600 ng/μl). This mixture was transferred to an electroporation cuvette (Harvard Apparatus) with a 1 mm gap electrode. Using a BTX ECM830 Square-pulse Electroporator (Harvard Apparatus), embryos were electroporated with six 3 ms pulses of 30V in 100 ms intervals. Electroporated zygotes were immediately implanted in pseudo-pregnant dams via oviduct transfer, designating the transfer/implantation date as embryonic day 0.5 (E0.5). Embryos were collected at E18.5, assessed for gross morphology defects, imaged under brightfield, and fixed in 4% paraformaldehyde.

Genomic DNA was extracted from embryonic tail tips and the respective editing sites in Exons 2 and 3 of *Zfp750* were PCR amplified using the following primers:

| Zfp750-Ex2-GT-F | AAAGAGGAGGCAGCAGTGAC |
| --- | --- |
| Zfp750-Ex2-GT-R | GCGTGTGGAATGCAGACTTG |
| Zfp750-Ex3-GT-F | CTGCCACTCAATCTGTCCGT |
| Zfp750-Ex3-GT-R | GCCAAGGCAGGAAGATCACA |

PCR products were then analyzed by Sanger sequencing. Sequence traces were first manually curated for preliminary scoring of homology-directed repair (HDR) and non-homologous end joining (NHEJ) events. Sequences were also analyzed using the Inference of CRISPR Editing (ICE) deconvolution tool from Synthego. ([https://ice.synthego.com](https://ice.synthego.com/); Synthego Performance Analysis, ICE Analysis. 2019. v2.0.)

*Generation of mutant alleles for stable mouse lines*

To generate *Zfp750* p.Ser154Pro knock-in (*Zfp750^S154P^*) and Exon2 frame-shift mice, C57BL/6NJ (JR 5304) mouse zygotes were electroporated with Cas9 ribonucleoprotein complex and ssDNA donor oligo for the *Zfp750^S154P^* mutation. Electroporated zygotes were surgically transferred to pseudo pregnant recipient females and allowed to come to term. Subsequent founder mice were genotyped for CRISPR editing as described above. Founders carrying the p.Ser154Pro were selected and bred to homozygosity. Founders harboring a +1 insertion (G at chr11 121513693) were also selected and crossed to C57BL/6NJ mice for maintenance as heterozygotes.

*Timed Matings and Embryonic Harvests*

Mice heterozygous for the Exon 2 +1 insertion mutation were crossed to generate -/- and +/+ embryos for phenotypic analyses. Embryos were collected at embryonic (E) day E14.5 and E15.5, and limbs snap-frozen in liquid nitrogen for subsequent RNA extraction. Embryonic samples were harvested at E13.5, 14.5, and 15.5 for histology and immunofluorescence. Embryonic limbs were also collected at E15.5 for scanning electron microscopy (SEM). Embryos were harvested at E18.5 for gross phenotype and Toluidine Blue barrier function assay analyses. Wild-type and mutant embryos were littermates and therefore of approximately the same gestational age.

*RNA-sequencing and data analysis*

Murine embryos were harvested at E15.5. Embryonic paws were dissected and immediately flash frozen in liquid nitrogen and stored at -80˚C. RNA was extracted from right paws of homozygous *Zfp750* mutant and wild-type embryos (N = 4) using RNAqueous™ Total RNA Isolation Kit (Invitrogen; cat# AM1912). Zebrafish injected with control or *znf750* morpholino (~40 embryos per replicate, three replicates) were used for extracting total RNA. Total RNA samples were sent to Novogene Inc. for mRNA-sequencing (RNA-seq) following the standard Illumina protocol. Three biological replicates is the standard minimum for most biological experiments, balancing between cost and the ability to calculate a mean and standard deviation.

RNA-seq raw reads were adapter-trimmed and quality filtered using TrimGalore (<https://github.com/FelixKrueger/TrimGalore>) and checked using fastQC tool (<https://www.bioinformatics.babraham.ac.uk/projects/fastqc/>). Filtered reads were mapped on mm39 or danRer11 reference genome using STAR (Dobin et al. 2013). Reads mapped to the genes were quantified with featureCounts (Liao et al. 2014) and differential gene expression analysis was performed using DESeq2 (Love et al. 2014). DEQ2 software applies a generalized linear model framework that is analogous to ANOVA. It uses the *Wald Test* and the *Likely Ratio Test* to detect differentially expressed genes.

*Histology and immunofluorescence*

Murine embryos were harvested at E13.5, E14.5, and E15.5 fixed in 4% paraformaldehyde overnight, then subsequently stored in 70% ethanol. Heads and limbs were further processed and embedded in paraffin. For histological analysis, serial coronal paraffin sections (7 µm thick) were stained with hematoxylin and eosin. Images were acquired using a Nikon Eclipse E-600 (Nikon, Melville NY). Oral adhesions were scored on coronal histological sections by evaluating the presence of contacts between the mandibular, maxillary, and lingual epithelium. A total of 39 Zfp750 frameshift embryos (22 *Zfp750^fs/fs^*, 16 *Zfp750^+/+^*, 1 *Zfp750^fs/+^*) and 11 *Zfp750^S154P^* embryos (5 *Zfp750^S154P/S154P^*; 6 *Zfp750^+/+^*) were analyzed in a blinded manner.

For immunofluorescent staining, 4% PFA fixed, paraffin embedded slides were de-paraffinized and then treated with 0.05% Tween-20 citric acid antigen retrieval as previously described (Paul et al. 2017). Samples were blocked with 3% normal goat serum (Vector Laboratories, Burlingame, CA) for 1 hour at RT, incubated in primary antibody for 1 hour at RT, followed by PBS washes before being incubated in secondary antibody for 1 hour at RT. Nuclear DNA was labeled with Hoechst 33342 (1/10,000). Samples were mounted using ProLong^TM^ Diamond Antifade Mountant. Images were acquired with a Zeiss 980 confocal microscope (Zeiss, Oberkochen, Germany) and processed using Fiji software.

*Scanning Electron Microscopy*

E15.5 murine embryos were fixed in 2.5% glutaraldehyde in 0.1M Na-cacodylate buffer and stored in fixative solution at 4˚C for no more than a week before scanning using a Hitachi S-4800 microscope.

*Barrier Function Assay*

Murine embryos were harvested at E18.5 for skin barrier function assessment via dye exclusion assay using toluidine blue. Embryos (6 embryos per genotype from 2 separate litters) were euthanized by ex-sanguination, then placed through a graded series of methanol in water (25%, 50%, 75%, 100%, 2 minutes each at 4˚C). Embryos were then stained in 0.1% toluidine blue in water for 2 minutes on ice.

*Statistical analyses*

Significance was determined using appropriate statistical tests described in respective figure legends.

**Appendix Figures**

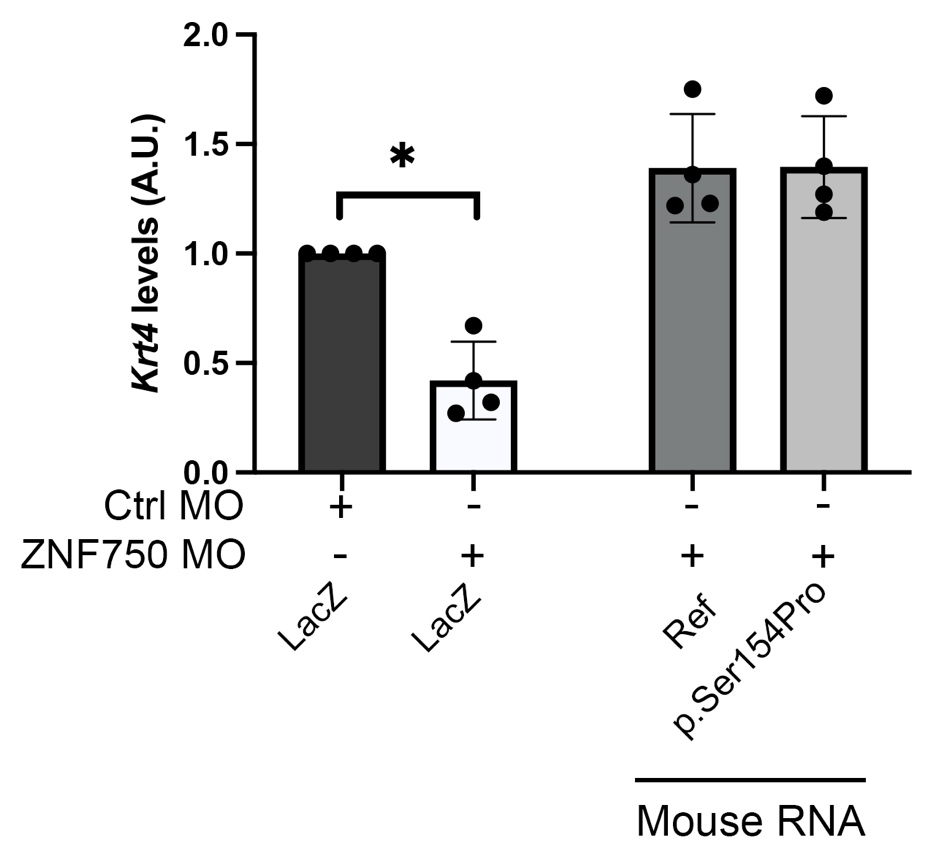

**Appendix Figure 1: Orthologous murine variant of *ZNF750* p.Ser160Pro (*Zfp750* p.Ser154Pro) does not affect *krt4* levels in zebrafish functional assays.** Levels of *krt4* expression are presented as a read out for functional validation of orthologous murine *zfp750* variants in a zebrafish assay. Zebrafish embryos (5.5 hpf) were injected individually with standard control (Ctrl) or *znf750* morpholinos (MO), then injected with: LacZ; mouse RNA encoding reference sequence; or *Zfp750* p.Ser154Pro variant. Significant differences in expression are denoted following One-way ANOVA; ∗ *P* < 0.05. Only significant differences are shown. Data are means of 4 biological replicates ± standard deviations.

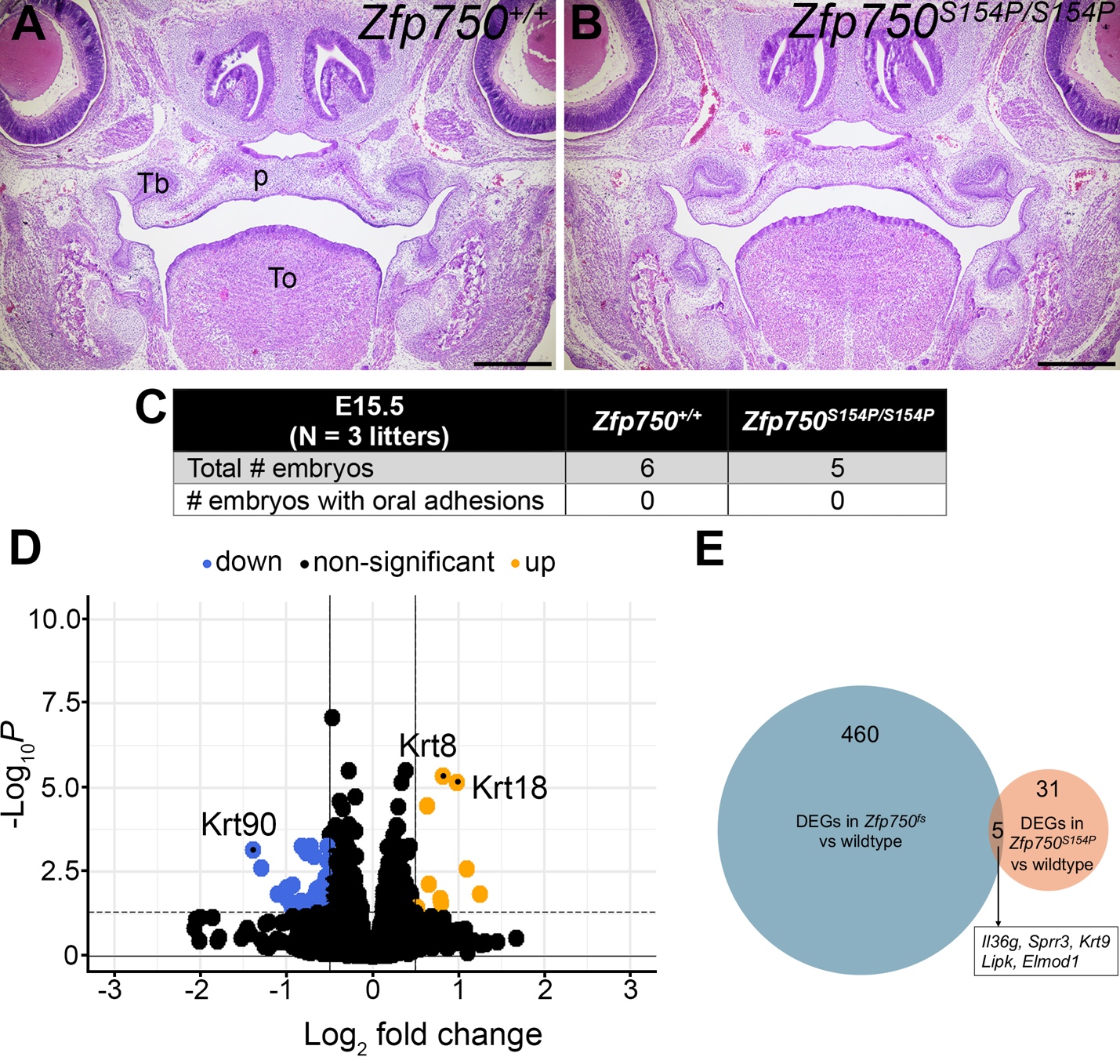

**Appendix Figure 2: *Zfp750^S154P^* exhibit proper palatogenesis and very few differentially regulated genes.** (**A-B**) Hematoxylin and eosin-stained coronal sections of *Zfp750^+/+^* (A) and *Zfp750^S154P/S154P^* E15.5 embryos (B). (**C**) Quantification of oral adhesions. (**D**) Volcano plot showing log2 fold change (x-axis) in gene expression between *Zfp750^S154P^* murine paw and corresponding wild-type control at E15.5. Adjusted *P*-values from the comparisons are plotted on y-axis. Up-regulated, down-regulated, and non-significantly regulated gene expression are shown as orange, blue, and black dots respectively. (**E**) Venn-diagram showing overlap between differentially regulated genes (DEGs) in E15.5 *Zfp750^fs^* vs. *Zfp750^S154P^*. To = tongue; Tb = tooth bud; p = palate. Scale bars = 500 µm

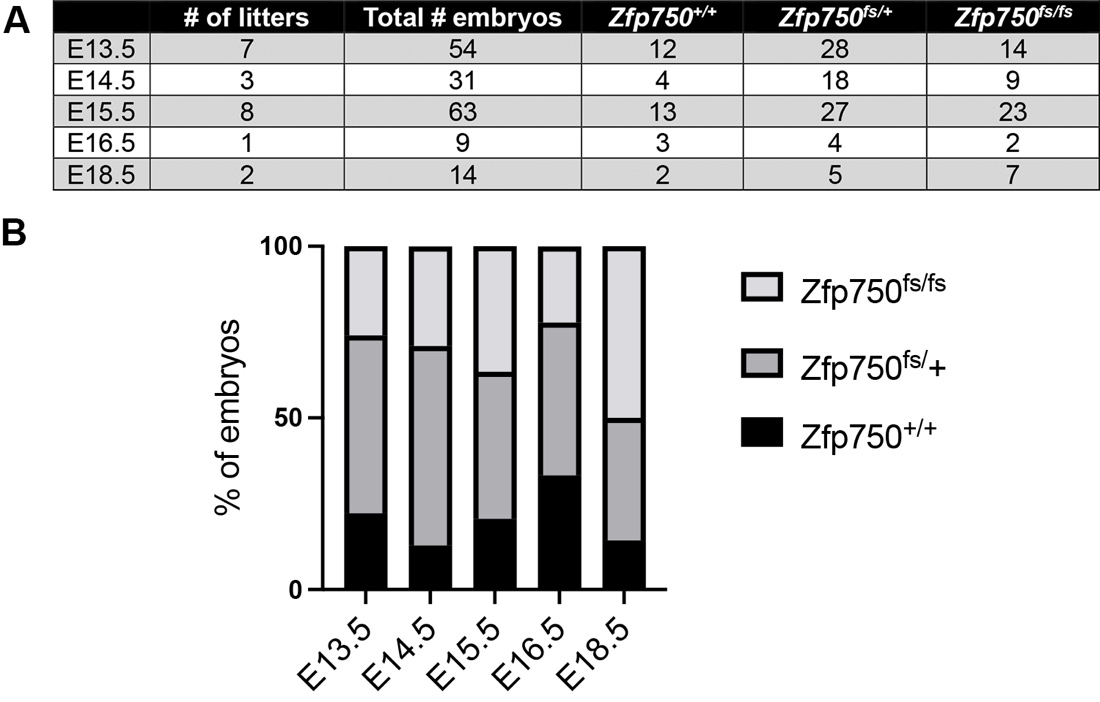

**Appendix Figure 3: *Zfp750^fs^* exhibit Mendelian ratios.** (**A**) Table summary of embryos collected at different developmental timepoints. (**B**) Graphical representation of data from the table. The proportion of embryos is not deviating significantly from the expected Mendelian ratios (25% for wildtypes and homozygotes, 50% for heterozygotes) following Chi-Square analysis.

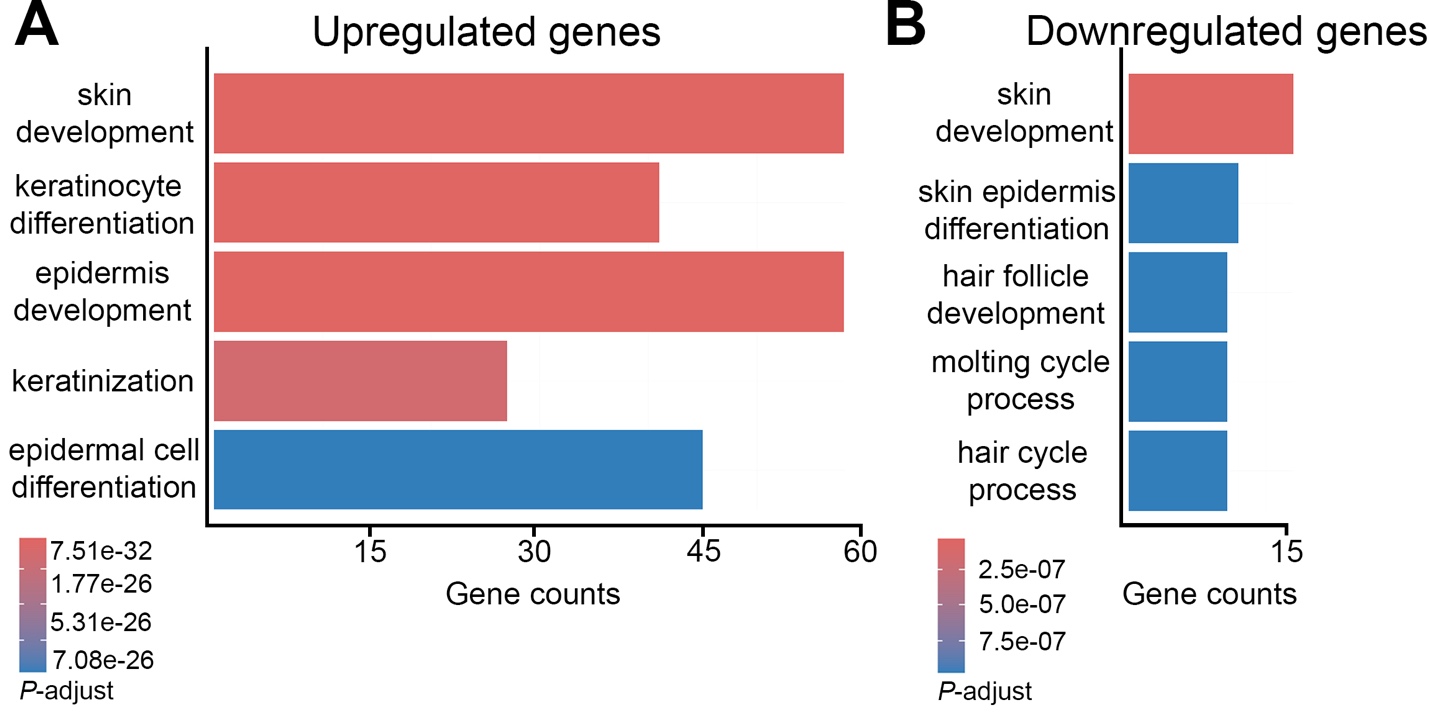

**Appendix Figure 4: Gene ontology analysis of RNA-seq from E15.5 forelimbs.** GO-term analysis of RNA-seq comparing E15.5 forelimbs from *Zfp750^+/+^* and *Zfp750^fs/fs^* embryos. (**A**) Upregulated genes. (**B**) Downregulated genes. Note the higher count for upregulated genes compared to the ones that are downregulated.

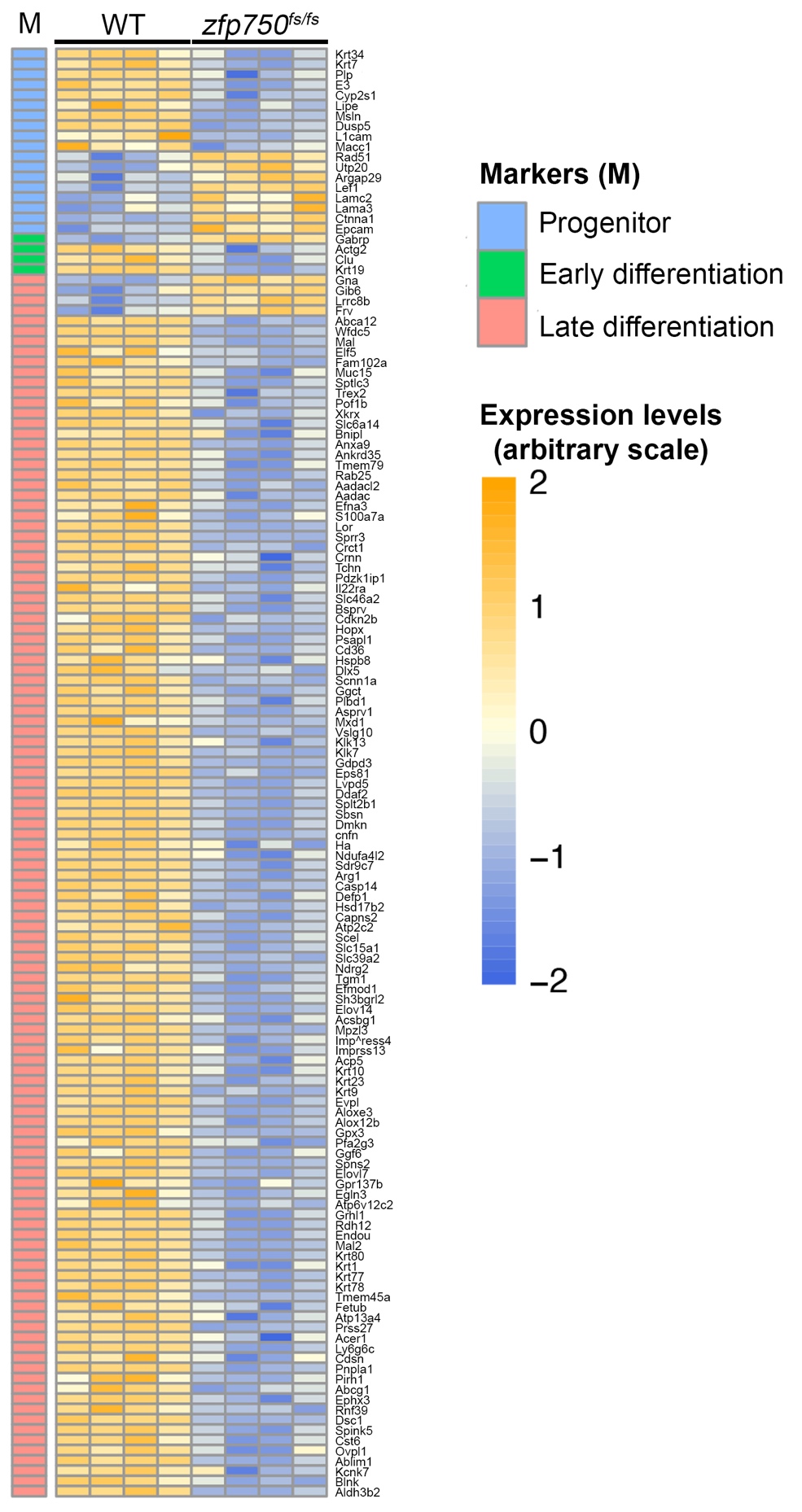

**Appendix Figure 5: Differentially expressed genes (larger view with gene list)**. This panel is the same as in Figure 5B with names of genes listed.

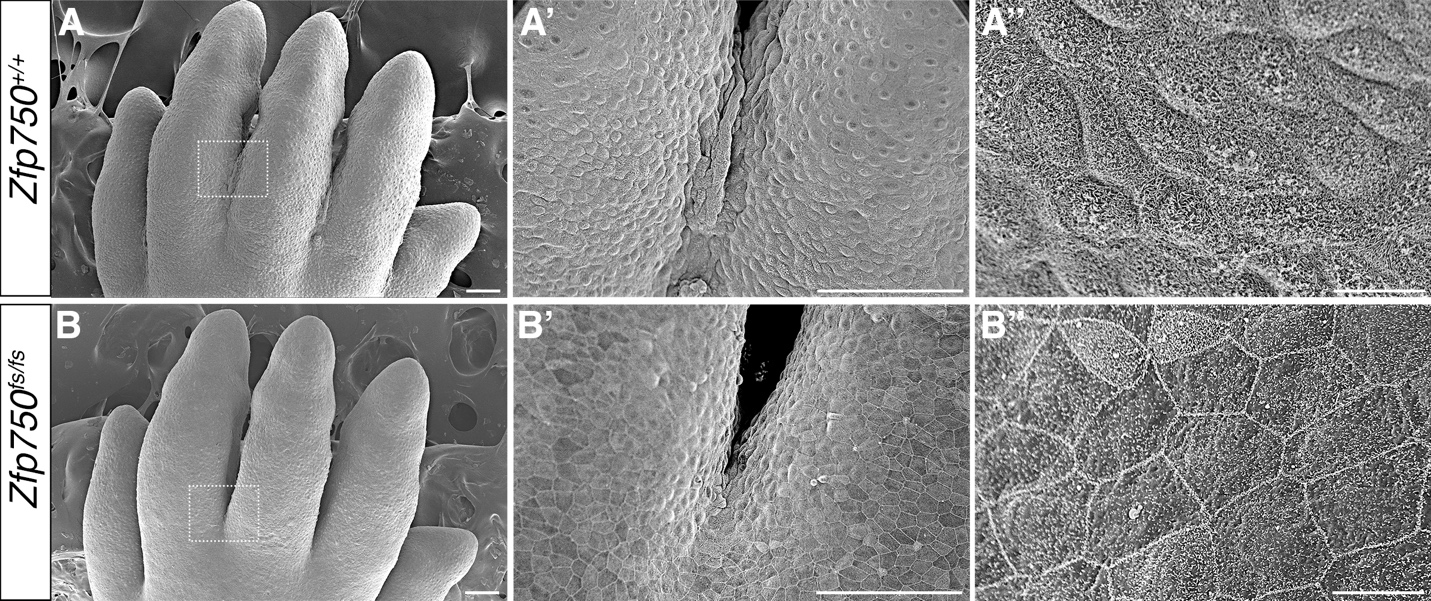

**Appendix Figure 6: *Zfp750^fs/fs^* leads to absence of interdigital ridge despite the presence of periderm-associated proteins**. (**A-B”**) Scanning electron microscopy of E15.5 hindlimbs reveal the presence of interdigital ridge (A [zoom-in of white boxes in A’]) and epidermal microvilli on the surface of wild-type (B”) but not in *Zfp750^fs/fs^* embryos (B, [zoom-in of white boxes in B’], B”). Scale bars: A-B’ = 100 µm; A”, B” = 10 µm

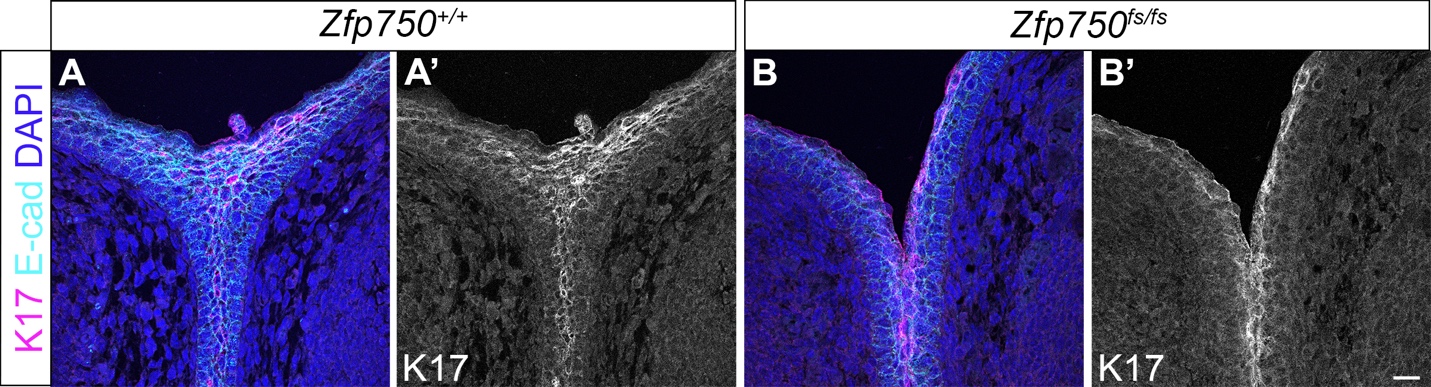

**Appendix Figure 7: Keratin 17 is detected in superficial cells of interdigital ridge in *Zfp750^fs/fs^***. (**A-B’**) Immunofluorescent dual staining of transverse digital section of E14.5 wildtype (A, A’) or *Zfp750^fs/fs^* (B, B’) for Keratin 17 and E-cadherin (Keratin 17 single channel shown in black and white). Scale bar = 20 µm

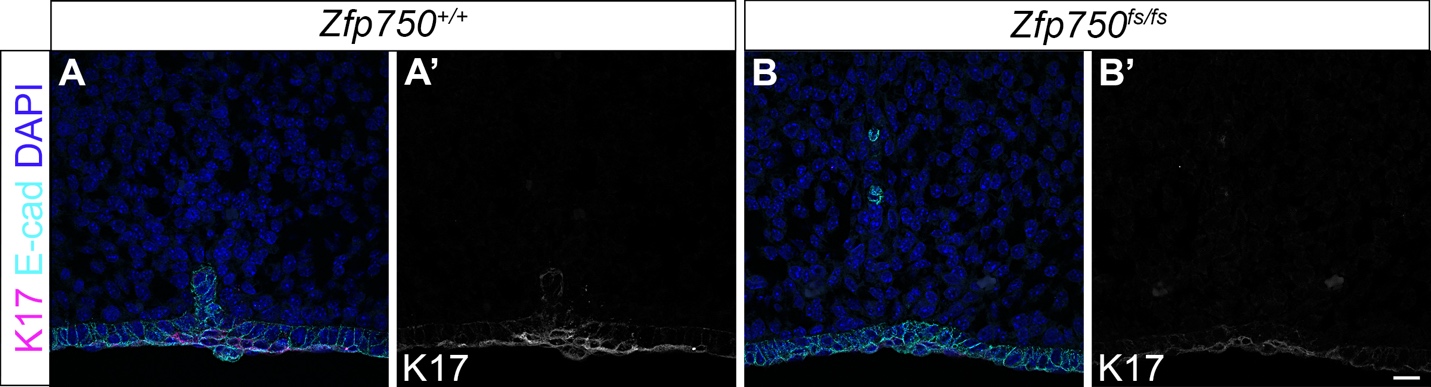

**Appendix Figure 8: Keratin 17 is detected in superficial cells of fused palatal shelves in *Zfp750^fs/fs^***. (**A-B’**) Immunofluorescent dual staining of coronal section of E14.5 wildtype (A, A’) or *Zfp750^fs/fs^* (B, B’) for Keratin 17 and E-cadherin (Keratin 17 single channel shown in black and white). Scale bar = 20 µm

**Appendix Files**

**Appendix Table 1:** Details of ZNF750 variants detected in OFC patients

**Appendix Table 2**: RNA-seq of zebrafish embryos at 5.5 hpf injected with *znf750* MO or with standard control MO.

**Appendix Table 3**: RNA-seq of paws harvested from mouse embryos at E15.5 that are *Zfp750^S154P/ S154^* or wildtype

**Appendix Table 4**: RNA-seq of paw harvested from mouse embryos at E15.5 that are *Zfp75 ^fs/fs^* or wildtype
